# Determinants of Gastrointestinal Group B *Streptococcus* Carriage in Adults

**DOI:** 10.1101/2023.08.17.553755

**Authors:** Elise S. Cowley, Ibrahim Zuniga Chaves, Fauzia Osman, Garret Suen, Karthik Anantharaman, Andrew J. Hryckowian

**Author notes:** Address correspondence to Andrew Hryckowian.

## Abstract

**Background:** *Streptococcus agalactiae* (Group B *Streptococcus*, GBS) is a commensal Gram-positive bacterium found in the human gastrointestinal and urogenital tracts. Much of what is known about GBS relates to the diseases it causes in pregnant people and neonates. However, GBS is a common cause of disease in the general population with 90% of GBS mortality occurring in non-pregnant people. There are limited data about the predisposing factors for GBS and the reservoirs in the body. To gain an understanding of the determinants of gastrointestinal GBS carriage, we used stool samples and associated metadata to determine the prevalence and abundance of GBS in the gut microbiome of adults and find risk factors for GBS status.

**Methods:** We used 754 stool samples collected from adults in Wisconsin from 2016-2017 to test for the prevalence and abundance of GBS using a Taqman probe-based qPCR assay targeting two GBS-specific genes: *cfp* and *sip.* We compared the microbiome compositions of the stool samples by GBS status using 16S rRNA analysis. We compared associations with GBS status and 557 survey variables collected during sample acquisition (demographics, diet, overall health, and reproductive health) using univariate and multivariate analyses.

**Results:** We found 137/754 (18%) of participants had detectable GBS in their stool samples with a median abundance of 104 copies per nanogram of starting DNA. There was no difference in GBS status or abundance based on gender. Beta-diversity, Bray-Curtis and Unweighted UniFrac, was significantly different based on carrier status of the participant. Prior to p-value correction, 59/557 (10.6%) survey variables were significantly associated with GBS carrier status and 11/547 (2.0%) variables were significantly associated with abundance (p-value<0.05). After p-value correction, 2/547 (0.4%) variables were associated with GBS abundance: an increased abundance of GBS was associated with a decreased frequency since last dental checkup (p<0.001) and last dental cleaning (p<0.001). Increased GBS abundance was significantly associated with increased frequency of iron consumption (p=0.007) after p-value correction in multivariate models.

**Conclusions:** GBS is found in stool samples from adults in Wisconsin at similar frequencies as pregnant individuals screened with rectovaginal swabs. We did not find associations between risk factors historically associated with GBS in pregnant people, suggesting that risk factors for GBS carriage in pregnancy may differ from those in the general population. We found that frequency of iron consumption and dental hygiene are risk factors for GBS carriage in Wisconsin adults. Given that these variables were not assayed in previous GBS surveys, it is possible they also influence carriage in pregnant people. Taken together, this work serves as a foundation for future work in developing approaches to decrease GBS abundance in carriers.

## BACKGROUND

*Streptococcus agalactiae*, also known as Group B *Streptococcus* (GBS), is a Gram-positive commensal bacterium in the gastrointestinal (GI) and urogenital (UG) tracts of humans [1,2]. While GBS typically colonizes asymptomatically at these body sites, it can cause illnesses such as bacteremia, pneumonia, meningitis, and soft tissue infections especially in adults and children with co-morbidities [3–5].

Much of what is known about GBS is related to diseases in pregnant individual, fetuses, and infants. GBS colonization rates of pregnant people as detected by rectovaginal swabs range from 10-35%, globally, with rates in the United States ranging from 10-30% [6,7]. Based on selective culturing of rectovaginal swabs and urine samples, the risk factors for GBS colonization are history of tobacco use, hypertension, black race, and younger age [8]. This colonization can result in urinary tract infections, chorioamnionitis, post-partum endometritis, and bacteremia in pregnant people [9,10]. Invasive disease has been associated with pregnancy loss, stillbirth, and preterm delivery [11]. GBS can vertically transmit to the neonate during vaginal delivery or infect *in utero*, where it causes early onset GBS disease (0-6 days of life) and is a risk factor for late onset GBS disease (7-89 days of life), which are the leading causes of neonatal sepsis and meningitis [12–16]. In these contexts, GBS causes over 400,000 symptomatic maternal, fetal, and infant cases globally per year [11]. Historically, it has been accepted that the GI tract is the reservoir for GBS leading to vaginal colonization in pregnant people and work in nonpregnant females has shown that rectal colonization is a strong predictor of vaginal colonization [17,18].

While much of what is known about GBS and human disease is in the context of pregnancy, GBS presents a significant disease burden in non-pregnant adults by causing bacteremia, sepsis, and soft tissue infections [3–5]. GBS disease in nonpregnant adults has steadily increased from 3.6 cases per 100,000 in 1990 to 7.3 in 2007 and 10.9 cases per 100,000 in 2016 with even higher incidences observed among those over the age of 65 years old [3,19]. These clinical data support that the current GBS disease burden is predominantly in non-pregnant people, which is likely due to widely implemented prophylactic strategies to reduce pregnancy-related transmission of GBS to neonates [20–22].

Risk factors for GBS disease in the general population include obesity, diabetes, increased age, and black race [3,23]. Despite this morbidity and mortality, little is known about the factors that dictate GBS colonization. GBS has been documented in the GI and UG tracts of non-pregnant adults, where between 9-32% of healthy adults have detectable GBS via oral swab, urine sample, or anorectal swab [24–28]. Although GBS can be found at these body sites, it is unclear what the risk factors are for colonization at these body sites in the general population. Taken together with the diseases caused by GBS in pregnant people, fetuses, and neonates, there is a need to better understand the reservoirs of GBS and risk factors for its asymptomatic carriage, which can inform our understanding of invasive disease and transmission to at-risk patient populations. Considering the profound consequences of GBS infections in pregnant patients and neonates, characterizing the presence of GBS in the general population will aid in our understanding of a possible reservoir and mode of transmission of GBS to pregnant patients.

To gain an understanding of the determinants of gastrointestinal GBS carriage in the general adult population, we used stool samples collected by Survey of Health of Wisconsin (SHOW) for the Winning the War on Antibiotic Resistance (WARRIOR) project to determine the frequency and prevalence of GBS in the gut microbiomes of adults in Wisconsin [29]. The samples from the WARRIOR study are from a representative cross section of adults in Wisconsin. The study collected extensive data from the participants on demographics, health, and diet. Understanding the prevalence of GBS in the general population and factors that may be associated with a higher abundance of GBS will further our understanding of the pathophysiology of GBS and inform new therapeutic options. Using samples from the general population will help us identify risk factors for GBS carriage that can inform our understanding of GBS ecology and help guide alternative approaches to coping with this pathogen in a variety of at-risk human populations.

## METHODS

### Stool samples

Human stool samples were previously collected and banked by SHOW from 2016-2017 as part of the WARRIOR project, which was reviewed and approved by the University of Wisconsin-Madison Institutional Review Board (Protocol #2013-0251) [29,30]. For our study, we used 754 WARRIOR stool samples from participants who agreed to have their samples used in future research. Our study was reviewed and approved by the University of Wisconsin-Madison Institutional Review Board (Protocol #2021-0025).

### DNA extractions from stool

Two methods were used to extract DNA from the stool samples. A subset of samples (455/754, 60%) had DNA remaining from a previous extraction and were used in this study [29]. Briefly, DNA was extracted using a bead-beating protocol with additional enzymatic lysis containing mutanolysin, lysostaphin, and lysozyme to help lyse Gram-positive bacterial cell walls. For the remaining 299 stool samples, in order to complete extractions in a high throughput manner, we extracted DNA using the DNeasy PowerSoil Pro kit (Qiagen) following the manufacturer’s instructions, starting with 50-250 mg stool and using a TissueLyzer II at 4°C for the homogenization followed by elution of DNA in 75 μL of solution C6.

### DNA extractions from bacterial cultures

For positive controls for quantitative PCR (qPCR) assays, we grew three strains of *S. agalactiae* in tryptic soy broth (Neogen NCM0004A) aerobically overnight at 37°C, including: *S.agalactiae* COH1, *S. agalactiae* 10/84, and *S. agalactiae* A909 (obtained from Katy Patras, Baylor College of Medicine). For a negative control, we grew *Clostridioides difficile* 630 in brain heart infusion broth (Neogen NCM0016A) anaerobically overnight at 37°C. One milliliter of each overnight culture was pelleted and the supernatants were removed. The DNA from the remaining cell pellets were extracted using the DNeasy Blood and Tissue kit (Qiagen) according to the manufacturer’s instructions for Gram-positive bacteria.

### DNA quantification

All DNA used in qPCR assays was quantified in duplicate on 96 well plates using the Quant-iT double stranded DNA broad range kit (Invitrogen) with a standard curve from 0 ng/μL – 100 ng/μL DNA. Fluorescent signal was read with Synergy HTX plate reader with excitation of 485/20 and emission of 528/20 and an auto-gain setting. Final DNA quantities in the samples were determined against the standard curve and by averaging duplicate readings.

### qPCR to quantify GBS prevalence and abundance in stool samples

We developed a multiplexed Taqman-based qPCR approach targeting two GBS-specific genes, *cfb* and *sip.* These genes were previously used in qPCR protocols to identify the bacterium [31–34] and the assay we developed was based on a protocol previously created for *sip* [35].

For *sip,* we used the primers: 5’-CAG CAA CAA CGA TTG TTT CGC C-3’ and 5’-CTT CCT CTT TAG CTG CTG GAA C-3’, targeting a 171 base pair region. The Taqman probe for *sip* was: 5’-FAM-AGA CAT ATT -ZEN -CTT CTG CGC CAG CTT TG-3IAkfQ-3’. For *cfb,* we used the primers: 5’-GAA ACA TTG ATT GCC CAG C-3′ and 5′.-AGG AAG ATT TAT CGC ACC TG-3′, targeting a 99 base pair region. The Taqman probe for *cfb* was 5’-HEX-CCA TTT GAT AGA CGT TCG TGA AGA G-3BHQ-1 -3’. We ran each reaction in triplicate with both *sip* and *cfb* probes and included positive controls (DNA from three GBS strains), a negative control (DNA from *C. difficile*), and we created a standard curve for each gene from 5×10^-3^ ng/µL to 5×10^-11^ ng/µL using serial dilutions of synthetic copies of target genes (gBlocks, IDT). All reactions totaled 19.25 µL and included 5 µL of DNA (either extracted from cultures of bacteria, extracted from stool samples, or synthetic DNA for the standard curve), 7.5 µL TaqPath qPCR Master Mix, CG, 0.75 µL of a 10 µM stock each primer (1 reverse and 1 forward for each gene target for 4 total primers), 0.3 µL of a 10 µM stock of each probe, and 3.15 µL water. Samples were held at 96°C for 5 minutes followed by 50 cycles of 96°C for five seconds, 58°C for 10 seconds, and 72°C for 20 seconds [35]. All qPCR was performed on an Applied Biosystems QuantStudio 7 instrument.

We determined GBS status by comparing the threshold cycle of each sample to the negative control. Samples with threshold cycles below the negative control for both *sip* and *cfb* were considered negative for GBS. Samples with threshold cycles above the negative control for both genes were positive for GBS and the abundance was determined by comparing cycle threshold against the standard curve.

### 16S rRNA marker gene analysis

We used 16S rRNA marker gene data previously generated for the WARRIOR samples [29]. Briefly, the V4 region of the 16S rRNA gene was sequenced on an Illumina MiSeq using 2 × 250 paired-end reads at the University of Wisconsin Biotechnology Center. Negative controls were included during each step of extraction and amplified and sequenced with the same protocol as described above.

The resulting fastq files were processed with the software QIIME 2 v2021.4 [36]. Demultiplexed raw sequences were imported using the Casava 1.8 format and denoised using DADA2 [37] (via qiime-dada2 plugin) to generate a feature table containing amplicon sequence variants (ASV). ASVs were aligned with MAFFT to construct a phylogenetic tree with fasttree [38]. Taxonomy was assigned using the classify-sklearn naive Bayes taxonomy classifier [39] against the Silva_138 database for 16S rRNA genes [40]. A feature and a taxonomy table, together with the phylogenetic tree were imported in R 4.1.2 as a phyloseq object [41] for further analysis. A total of 756 samples, including 29 negative controls and 727 stool samples, were processed with R. Contaminationwas accounted for by eliminating features based on the prevalence of ASVs in the negative controls using the Decontam package [42], and by removing eukaryotic, chloroplast, mitochondrial or unassigned sequences. Finally, samples without GBS-associated data or with less than 5000 reads were removed. The resulting 693 samples were processed as follows.

For the alpha and beta diversity analysis, samples were rarefied to an even depth of 8396, corresponding to the minimum read count in the dataset. Alpha diversity metrics (Shannon, inverse Simpson, and total observed ASVs) metrics were calculated with Phyloseq. Beta diversity was used to quantify the dissimilarities between the samples. First, we calculated 3 different types of distance matrices (Bray-Curtis, weighted and unweighted UniFrac and then plotted the ordinations using a principal component analysis (PCoAs). PCoAs were plotted for GBS carrier status, and the logarithmic transformation of the average copy number of GBS. Moreover, to confirm our findings, we extracted the distances for each matrix and plotted the average distance between and within sample types. Average distances were plotted as box plots and significant differences based on the carrier status of GBS were tested using a one-way ANOVA and Tukey’s HSD post-hoc test.

#### Linear correlations with GBS prevalence

To evaluate associations with alpha diversity, simple linear regressions were calculated for prevalence and copy number against the Inverse Simpson’s, Observed features, and Shannon’s indices. For these analyses, we included only the samples where GBS was detectable via qPCR.

#### Differential taxonomic abundance

Differential taxonomic abundance was obtained using the ANCOM-BC package in R using default parameters and Benjamini-Hochberg procedure for P-value corrections [43]. Linear discriminant analysis effect size (Lefse) [44] was done using the R package microbiomeMarker [45]. Finally, the QCAT package was used to evaluate microbiome markers using copy numbers as it allows for continuous variables [46].

#### Random forest classification

To identify taxa that discriminate between the presence and absence of GBS, we used a random forest classifier algorithm from the random forest R package [47]. In summary, we trained and tested our model using the “out of bag” (OOB) error to estimate our model error. The number of trees was set to 1000 and only ASVs with relative abundancesL≥ 0.01% were included as input. The classifier was trained on a random selection of 70% of the database composed of 693 samples and 1099 ASVs and validated using the remaining 30%. Finally, prediction performance was measured by the OOB error rate and the mean decrease in Gini coefficient, a measure of how each variable contributes to the homogeneity of the nodes and leaves in the resulting random forest, for each ASV.

### Metadata variable acquisition and selection

We acquired participant metadata from SHOW, which biobanks the samples and manages the data repository associated with the samples. For our analysis of all samples, we selected relevant variables that fit into three categories: demographics, health, and diet. For participants who identified as female, we also analyzed data on reproductive health. Data were collected for some study participants across multiple years, we selected variables relevant for the years the stool samples were collected (2016-2017).

### Analysis of WARRIOR study metadata

#### Differences in GBS outcome by self-identified gender

To determine differences in carriage of GBS by self-identified gender, we used Pearson’s Chi-squared test with Yates’ continuity correction. To determine differences in abundance of GBS by self-identified gender and for those individuals with detectable GBS, we used Welch’s two sample t-test with unequal variances. We used R version 4.2.2 to run these comparisons.

#### Binary outcome

To determine associations between a binary outcome of GBS (presence or absence of GBS in the stool) and other predictors, we used a logistic regression model and reported odds ratios and confidence intervals. We conducted a univariate analysis first (each predictor in a bivariate analysis), then significant predictors from the independent models were used to construct three multivariate (adjusted) logistic models. Model 1 examined significant predictors from the univariate analysis, model 2 examined predictors commonly associated with GBS carriage or disease risk, as shown in the literature (African American, Type 2 Diabetes Mellitus, and smoking status)[8], and model 3 combined predictors from models 1 and 2. Odds ratio plots were created using log transforms of the odds ratios and 95% confidence intervals. Variables with missing data were addressed using the complete case analysis method. In this instance, any subject with no observations was removed from the specific analysis being conducted. After adjusting for missing data, a total of 555 variables were used for the binary outcome comparisons All analyses were conducted using STATA version 17 (StataCorp. 2021. *Stata Statistical Software: Release 17*. College Station, TX: StataCorp LLC).

#### Continuous outcome

Predictors associated with the abundance of GBS for participants with detectable GBS were examined using a linear regression model with coefficient estimates and 95% confidence intervals. We used a similar approach used above see <u>Binary outcome</u> above. Each predictor was independently tested and determined to either be fit for a multivariate model or not included in the model. We constructed the same three models specified above for the binary outcome. We constructed odds ratio plots as described above. Variables with less than 5% missing data were negligible and used as is, variables with up to 40% missing data were inputted using multiple imputation methods and variables with greater than 50% missing data were excluded from the analysis altogether. After adjusting for missing data, a total of 547 variables were used for the continuous outcome comparisons. All analyses were conducted using STATA version 17 (StataCorp. 2021. *Stata Statistical Software: Release 17*. College Station, TX: StataCorp LLC).

#### Subgroup analysis

A sub-group analysis in females was conducted using the same statistical tests for either the binary outcome or the continuous outcome with the same independent variables for each outcome. All analyses were conducted using STATA version 17 (StataCorp. 2021. *Stata Statistical Software: Release 17*. College Station, TX: StataCorp LLC).

#### Diagnostics and missing data

We ran collinearity diagnostics to identify independent variables with significant relations as this is disruptive to models. We used a variance inflation factor (VIF) to measure the tolerance of variance. Variables with a VIF value greater than 10 were removed from the models. Secondary pairwise correlations were conducted to supplement our decision to remove collinear variables. We corrected for multiple comparisons effect on the p-value using a false discovery rate (FDR) using the Benjamini-Hochberg (BH) procedure. All p-values were assigned a rank within the outcome group and a critical value using the BH procedure was calculated using false discovery rates of 5% and 10% to discover which worked best for the data. Critical values less than the significant original p-values were also regarded as significant. We graphed the odds from the multivariate analysis using coefficient plots with confidence intervals. All analyses were conducted using STATA version 17 (StataCorp. 2021. *Stata Statistical Software: Release 17*. College Station, TX: StataCorp LLC).

## RESULTS

### Participant Demographics

For this study, we used stool samples from 754 unique individuals across the state of Wisconsin, collected between 2016-2017. Our samples came from 432 (57%) self-identified females and 322 (43%) self-identified males with ages ranging from 18 to 57 (**Table 1**).

**Table 1.**
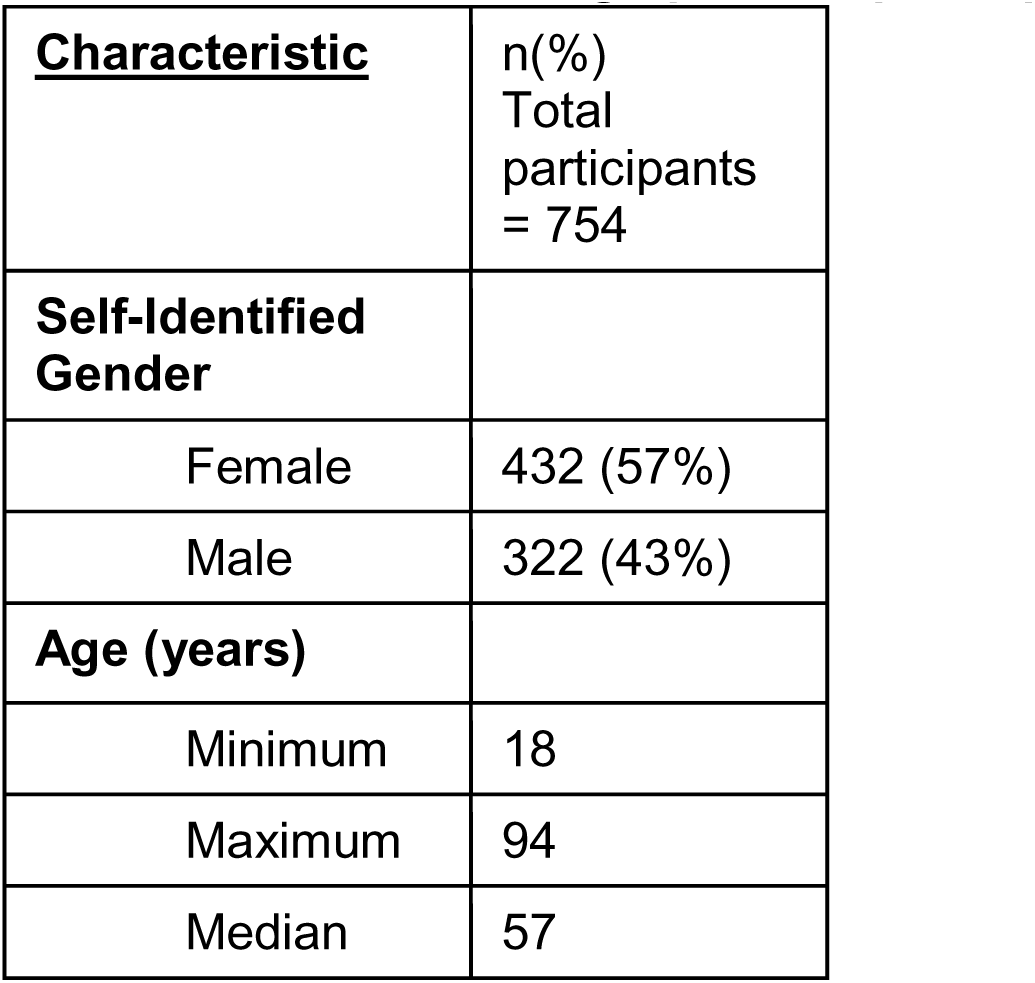
Baseline demographics of participants.

| **<u>Characteristic</u>** | n(%)<br>Total<br>participants<br>= 754 |
| --- | --- |
| **Self-Identified Gender** |  |
| Female | 432 (57%) |
| Male | 322 (43%) |
| **Age (years)** |  |
| Minimum | 18 |
| Maximum | 94 |
| Median | 57 |

### GBS is present in gut microbiomes of at widely variable abundances

Using a qPCR-based method, we found that 137/754 (18%) of all participants had detectable GBS in their stool samples with 79/432 (18%) of self-identified females and 58/322 (18%) of self-identified males having detectable GBS (**Figure 1A**). There was no significant difference in proportion of either gender based on GBS status. For samples with detectable GBS, we quantified the amount using a standard curve. The samples contained between 5 to 6,800,532 copies of GBS target DNA per starting nanogram of total DNA (**Figure 1B**). There was no difference in abundances of GBS in the stool based on the gender of the study participants.

**Figure 1.**
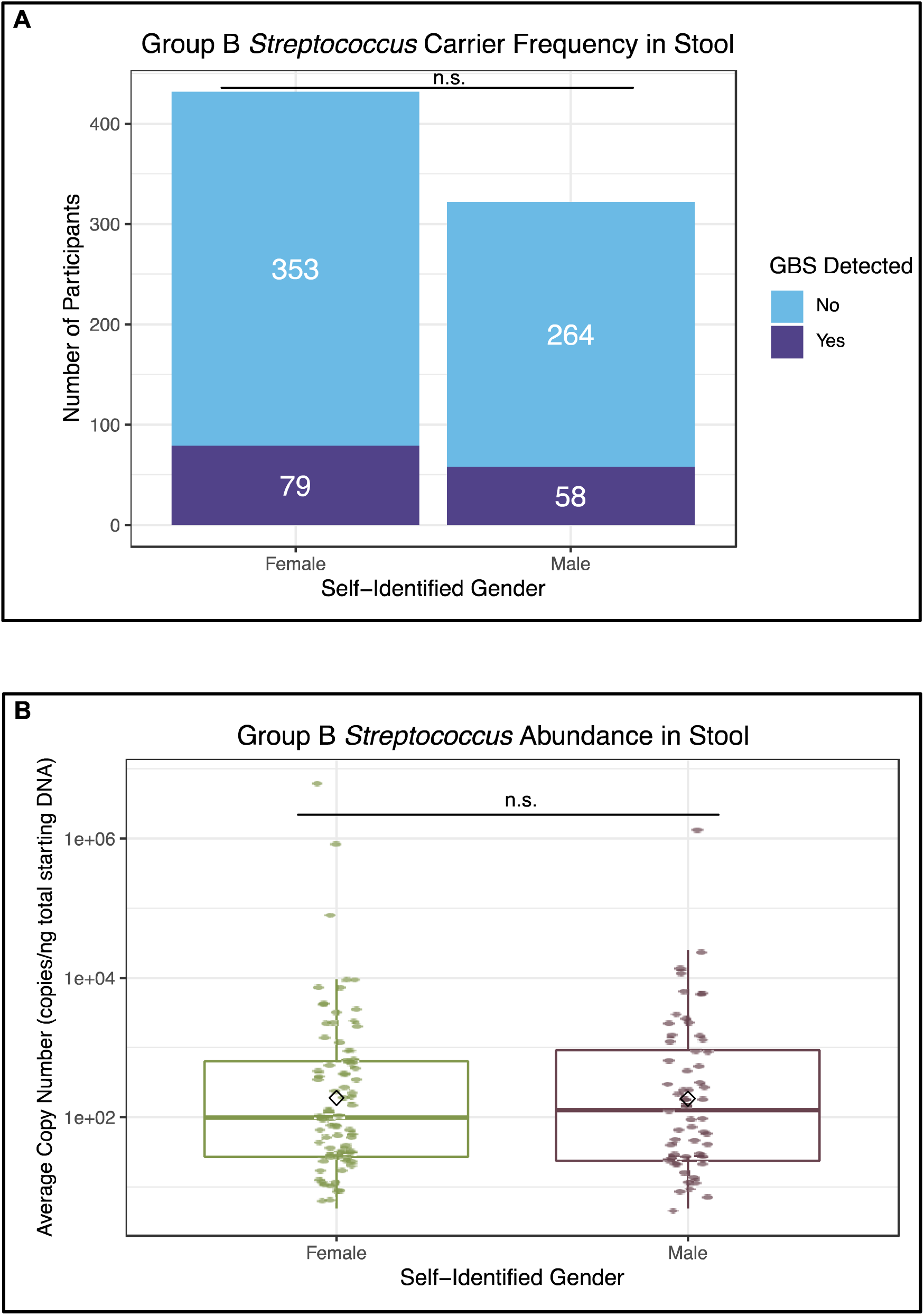
Carriage of GBS and abundance by self-identified gender. **A.** Presence or absence of GBS in stool samples as determined by qPCR. No significant difference was found in the prevalence of GBS between genders as indicated by “n.s.” as determined by Chi-squared tests. **B.** Abundance of GBS in stool samples with identifiable GBS as determined by qPCR. Each colored point represents GBS abundance in an individual sample, diamonds represent the mean copies of GBS for each gender and the box plots represent median, interquartile ranges of GBS abundance, and standard deviation for each gender. The median copy number of GBS per starting amount of DNA was 104 and the average was 66,856. The median copy number for self-identified females was 98, with a mean of 97,621, a standard deviation of 768,659 and an interquartile range (IQR) of 605. For males, the median copy number was 126, mean 24,951, standard deviation of 178,687, and an IQR of 898. No significant difference was found in GBS abundance among male and female carriers as indicated by “n.s.” as determined by Welch’s two sample t-test with unequal variances.

### Differences in gut microbiome composition of WARRIOR study participants based on GBS carrier status

Of the 754 samples, 693 (92%) had available 16S rRNA marker gene sequencing performed in a previous study [29]. In this subset of samples, no significant correlations were found between GBS presence or abundance and alpha diversity metrics (Inverse Simpson, observed ASV, or Shannon) using linear regression analysis (**Table 2**).

**Table 2.** Correlations between alpha and beta diversity metrics and GBS carriage or abundance.

| Diversity Metric | Presence or Absence p-value | Abundance p-value |
| --- | --- | --- |
| **Alpha Diversity** |  |  |
| Inverse Simpson | 0.449 | 0.999 |
| Observed ASV | 0.949 | 0.220 |
| Shannon | 0.785 | 0.636 |
| **Beta Diversity** |  |  |
| Bray-Curtis | 0.046* | 0.176 |
| Log transform |  | 0.013* |
| Weighted UniFrac | 0.205 | 0.810 |
| Log transform |  | 0.290 |
| Unweighted UniFrac | 0.049* | 0.183 |
| Log transform |  | 0.064 |

\*indicates p-value<0.05

To evaluate differences in bacterial communities we used a PCoA with different distance matrices (Bray-Curtis, Weighted and Unweighted UniFrac) grouped by GBS carriage and copy number (**Table 2**). Using a PERMANOVA test, we found that the bacterial communities were significantly different according to the presence/absence of GBS in PCoAs with the Bray-Curtis and Unweighted UniFrac distance matrices. Moreover, we found that bacterial communities were significantly different for the abundance of GBS with the log transform for the PCoAs with the Bray-Curtis dissimilarity matrix (**Table 2**).

Additionally, we determined differences in the average beta diversity distance by group based on GBS carrier status. We found statistically significant differences in the average Bray-Curtis, Weighted and Unweighted UniFrac distances across the three comparisons: intragroup distance between all samples with GBS compared to intragroup distance for samples without GBS (p<0.001 in all three indices), intergroup distance differences for samples with and without GBS compared to intragroup distances between samples without GBS (p<0.001 across all three indices), and finally intergroup distance differences for samples with and without GBS compared to intragroup distances between samples with GBS (p≤0.001 across all three indices) (**Figure 2**).

**Figure 2.**
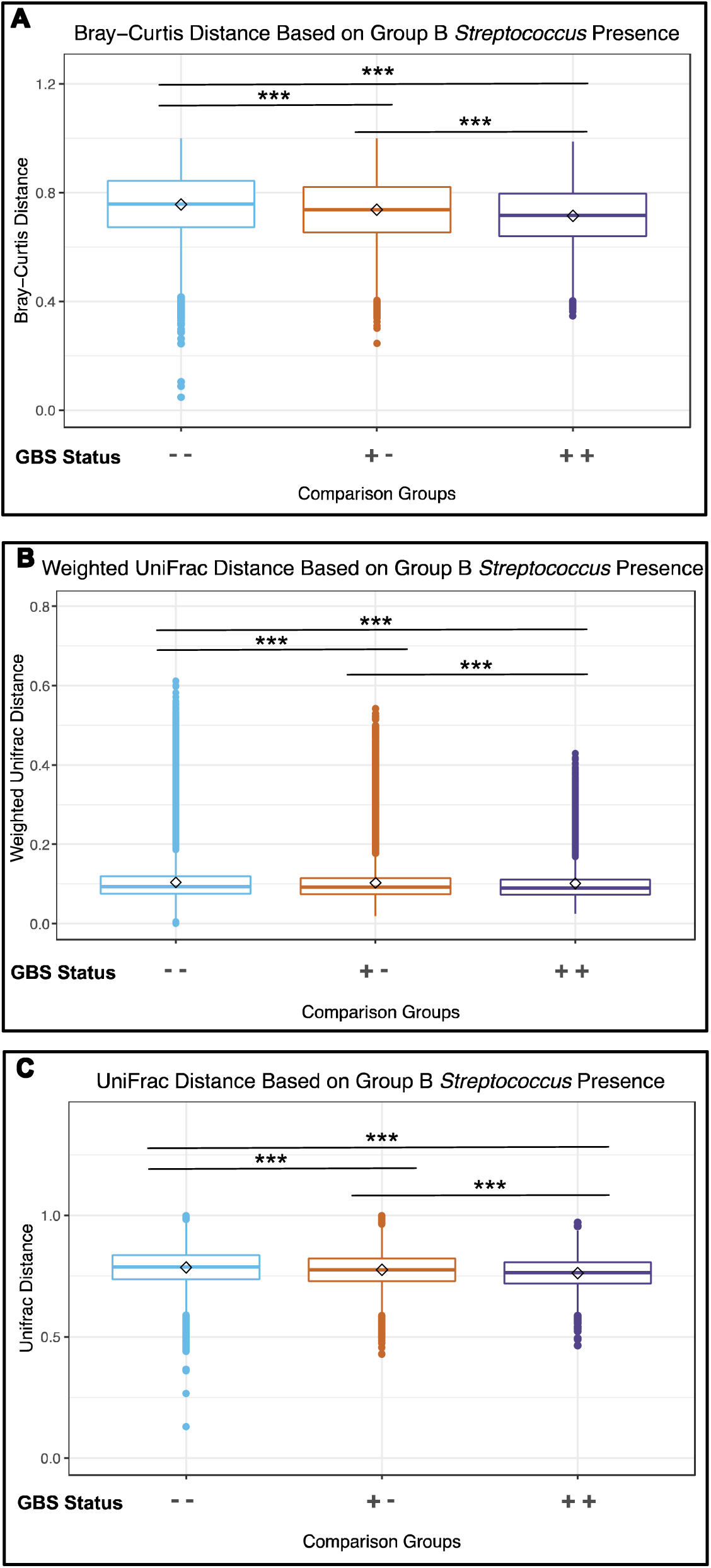
Beta diversity distance differences between GBS groups. Beta diversity distances for inter and intra group comparisons are plotted for three beta diversity indices. Plotted are the intragroup distances for samples without GBS (− −) intragroup distances for samples with GBS (+ +), and the intergroup distances for samples with and without GBS (+ −). The average distance for each of the three groups were compared to test for differences in beta diversity. The three beta diversity indices used were: **A.** Bray-Curtis, **B.** Weighted UniFrac, and **C.** UniFrac. The diamonds indicate the mean for each category. *** indicates a p-value ≤ 0.001.

We compared differential abundances of microbiome members at different taxonomic levels as a function of GBS carrier status. We found no differences at the phylum or ASV level, however at the genus level, we found that the relative abundance of two genera, *Ruminococcus* (p=0.0003 uncorrected, p=0.055 corrected) and *Monoglobus* (p=0.00057 uncorrected, p=0.057 corrected) trended towards being significantly differentially abundant when GBS is present, specifically both genera trended toward being more abundant when GBS is present.

To gain additional insights into microbes that differentiate GBS carriers from non-carriers, we ran random forest classifiers to identify community signatures predictive of GBS carrier status. This analysis could reliably predict GBS negative individuals, but not GBS positive individuals (error rates = 0.0035 and 0.992, respectively; **Supplemental Table 1**). Several taxa were identified through the random forest classification as good predictors of GBS status, including *Anaerostripes , Ruminococcus ,Collinsella, Dorea, Agathobacter, Blautia, Faecalibacterium, Bacteriodes, and Anaerostripes.* (**Supplemental Table 1**).

### Differences in characteristics of WARRIOR study participants based on GBS presence and abundance

We completed univariate and multivariate analyses on study participant metadata to understand which host characteristics were associated GBS carriage and which host characteristics were correlated with GBS abundance in carriers. For our analysis, we selected relevant variables that fit into three categories: demographics, health, and diet. For participants who identified as female, we also analyzed data on reproductive health which were not collected for participants who identified as male. The median and interquartile range for all variables considered in our multivariate statistical analysis are listed in **Supplemental Table 2**.

From our univariate analysis for carrier status (GBS present or absent in the stool), we found 59/557 (10.6%) variables were significantly associated with GBS carrier status (p-value < 0.05), but no variables were significantly associated after p-value corrections with FDR (**Supplemental Table 3**). Similarly, for our three multivariate models, we found no significant associations after p-value correction (**Figure 3, Supplemental Table 4**).

**Figure 3.**
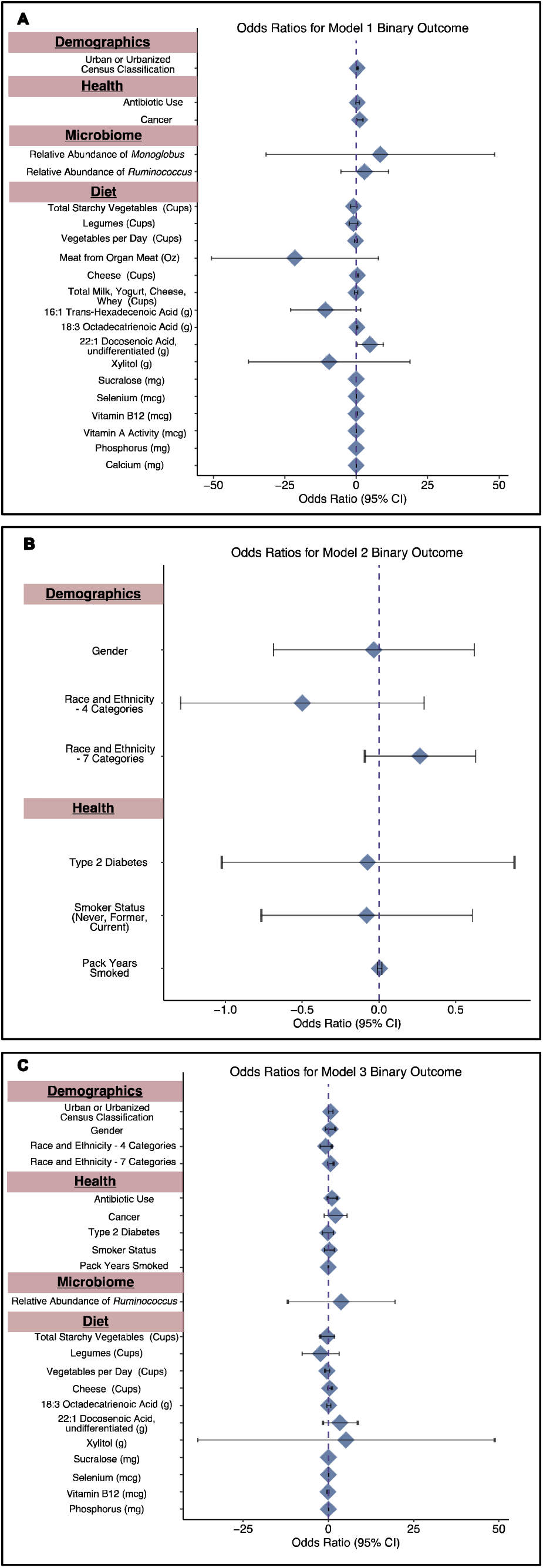
Odds ratios for multivariate models comparing presence or absence of GBS and participant characteristics. The diamonds indicate the odds ratios for each variable with error bars for the 95% confidence interval. **A.** Model 1 compared significant predictors from the univariate analysis prior to FDR correction with GBS presence or absence. **B.** Model 2 compared predictors commonly associated with GBS from previous studies and GBS presence or absence. **C.** Model 3 combined predictors from Models 1 and 2 with GBS carrier status. **indicates p-value ≤ 0.001 after FDR correction

For the univariate analysis of the abundance of GBS from participants with detectable GBS in their stool, we found 11/547 (2.0%) variables were significantly associated with GBS abundance (p-value < 0.05) prior to FDR correction. After FDR correction, we found 2/547 (0.4%) variables associated with GBS abundance: an increased abundance of GBS was associated with an increased time since last dental checkup (p<0.001) and last dental cleaning (p<0.001) (**Supplemental Table 5**). For the multivariate analysis, we found that in model 1, which included all the variables that were significant from the univariate analysis prior to p-value correction, higher GBS abundance was significantly associated with an increased frequency of iron consumption (p=0.007) after p-value correction. No other variables were significantly associated with GBS abundance in any of the other multivariate analyses (**Figure 4, Supplemental Table 6**).

**Figure 4.**
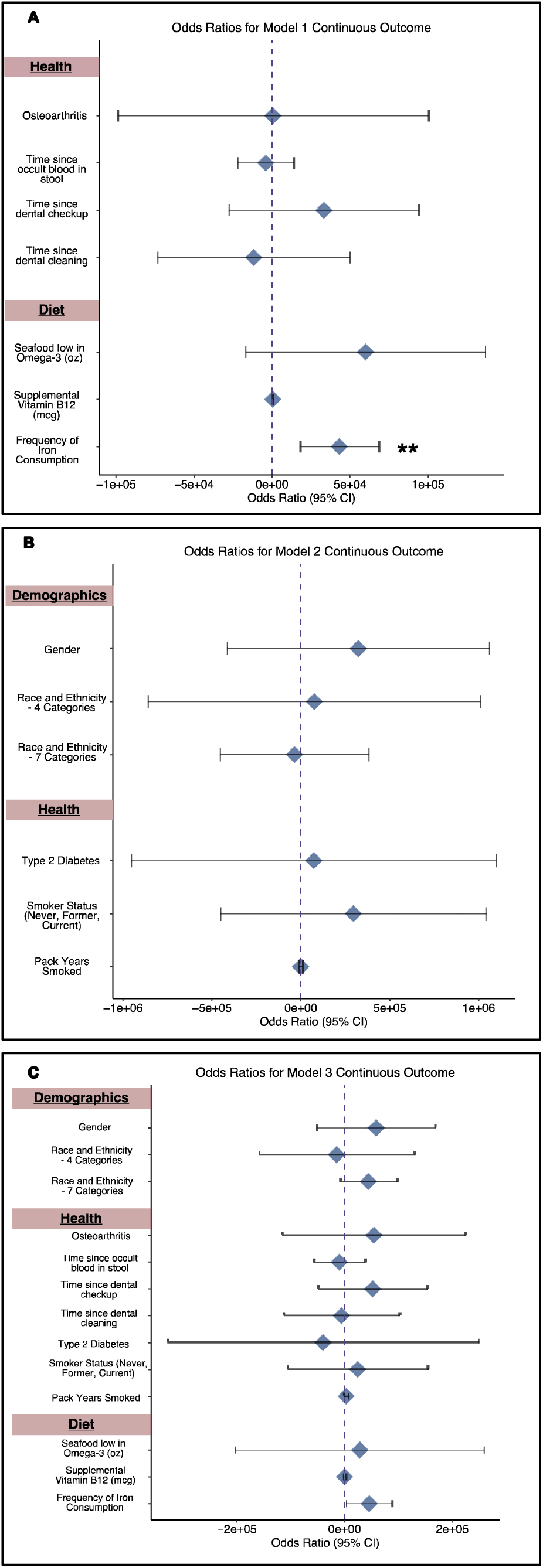
Odds ratios for multivariate models comparing GBS prevalence and participant characteristics. The diamonds indicate the odds ratios for each variable with error bars for the 95% confidence interval. **A.** Model 1 compared significant predictors from the univariate analysis prior to FDR correction with GBS prevalence. **B.** Model 2 compared predictors commonly associated with GBS from previous studies and GBS prevalence. **C.** Model 3 combined predictors from Models 1 and 2 with GBS prevalence. **indicates p-value <0.01 after FDR correction

### Subgroup Analysis of Associations of GBS for Self-Identified Females

Since GBS has historically been studied in pregnant people, we performed a univariate sub-group analysis of all self-identified females to investigate if there was a difference in the sub-population that were not present in the general population. Similar to the general population, in self-identified females, we tested for variables that differentiate GBS carrier status and for GBS abundance in the subset of female study participants with detectable GBS. For carrier status, we found 6/556 (1.1%) variables associated with GBS presence or absence prior to p-value correction and we did not find any significant associations after p-value correction with FDR (**Supplemental Table 7**). We found 10/497 (2.0%) variables associated with GBS abundance prior to p-value correction and 2/497 (0.40%) significantly associated after p-value correction (**Supplemental Table 8**). For abundance, similar to the analysis with all participants, an increased abundance of GBS was associated with a decreased frequency since last dental checkup (p<0.001) and last dental cleaning (p<0.001), after p-value correction (**Supplemental Table 8**). Unlike in the general population, we did not observe a correlation with GBS abundance and frequency of iron consumption.

## DISCUSSION

In this study, we used a biobank of 754 stool samples and associated data collected from adults in Wisconsin to better understand the host and microbiome-based factors that influence gastrointestinal GBS carriage.

We found that GBS is present in the stool from a representative cross-sectional sampling of adults in Wisconsin at rates like what is seen in pregnant individuals via rectovaginal swabs. We found that the abundance of GBS varied by orders of magnitude between individuals with GBS and there was no difference in the carrier frequency or abundance based on gender. These data indicate that GBS is common in the distal GI tracts of the general population, providing further evidence that the distal GI tract is an important reservoir for GBS in the human body [17,28].

We carried out 16S rRNA marker gene analysis of the samples in the context of GBS colonization and found that regardless of index used (Bray-Curtis, Weighted UniFrac, Unweighted UniFrac), beta-diversity was significantly different based on GBS carrier status. We did not find robust correlations between alpha-diversity or individual microbes that predicted GBS carriage or GBS abundance in individuals with GBS. It is possible that multiple microbiome configurations or functionally redundant microbiome members influence GBS carriage or that transient GBS carriage, as has been observed for pregnant people, obscures this assessment [48].

In addition, to identify host-centric variables that influence GBS carriage, we leveraged the extensive participant metadata collected as part of the WARRIOR study. While we did not find many statistically significant variables that correlated with GBS carrier frequency or abundance after p-value correction, prior to p-value correction, there were 59/557 (10.6%) variables for GBS carrier status and 11/547 (2.0%) for GBS abundance that were statistically significant (p<0.05). The two most robust associations we found after p-value corrections were between lack of dental care and higher iron consumption and increased GBS abundance. *Streptococci* are known inhabitants of the upper GI tract, including the oral cavity [28,49]. Our finding that a lack of dental care is associated with higher GBS burdens in the stool implies that poor dentition leads to an increase in *Streptococci* in the oral cavity, which can subsequently be observed more distal in the GI tract [50]. Iron is an essential nutrient for almost all living organisms, including bacteria, which have evolved strategies to scavenge environmental iron. These strategies enable bacteria to compete with other microbes within microbiomes and have been demonstrated to be important to overcome human immune defenses [51]. In particular, GBS encodes siderophores which aid in environmental iron acquisition [52] and elevated oral iron consumption may favor siderophore-dependent iron acquisition by GBS and increased GBS fitness in the distal GI tract. Alternatively, it is possible that iron indirectly impacts GBS fitness by influencing the abundance of microbes that compete or cooperate with it in the distal GI. Supportively, oral iron consumption has been previously shown to increase the amount of iron available to the gut microbiome and influence its composition [53,54].

Of note, we did not find associations between variables historically associated with GBS carriage in pregnant people in our study population. Specifically, neither GBS carriage nor GBS abundance in carriers was impacted by race, smoking status, age, or diabetes status (Figure 5). This challenges our current understanding of the risk factors for GBS carriage and highlights two possibilities for future study. First, it is possible that risk factors for GBS carriage in pregnancy are different than the risk factors in the general population. Second, it is possible that iron consumption and dental hygiene are risk factors for GBS carriage in pregnant people, but studies have not been done to address these connections. Therefore, further study of both pregnant and non-pregnant adults, informed by this work, will fill these important gaps in understanding relating to the life cycle of GBS. This understanding is a useful starting point developing new interventions, beyond antibiotics, to de-colonize GBS carriers and to prevent GBS infections in at-risk populations. These considerations are relevant given the rising incidence of antibiotic resistance in GBS and our emerging understanding of the collateral damage antibiotics have on human microbiomes [55–60].

**Figure 5.**
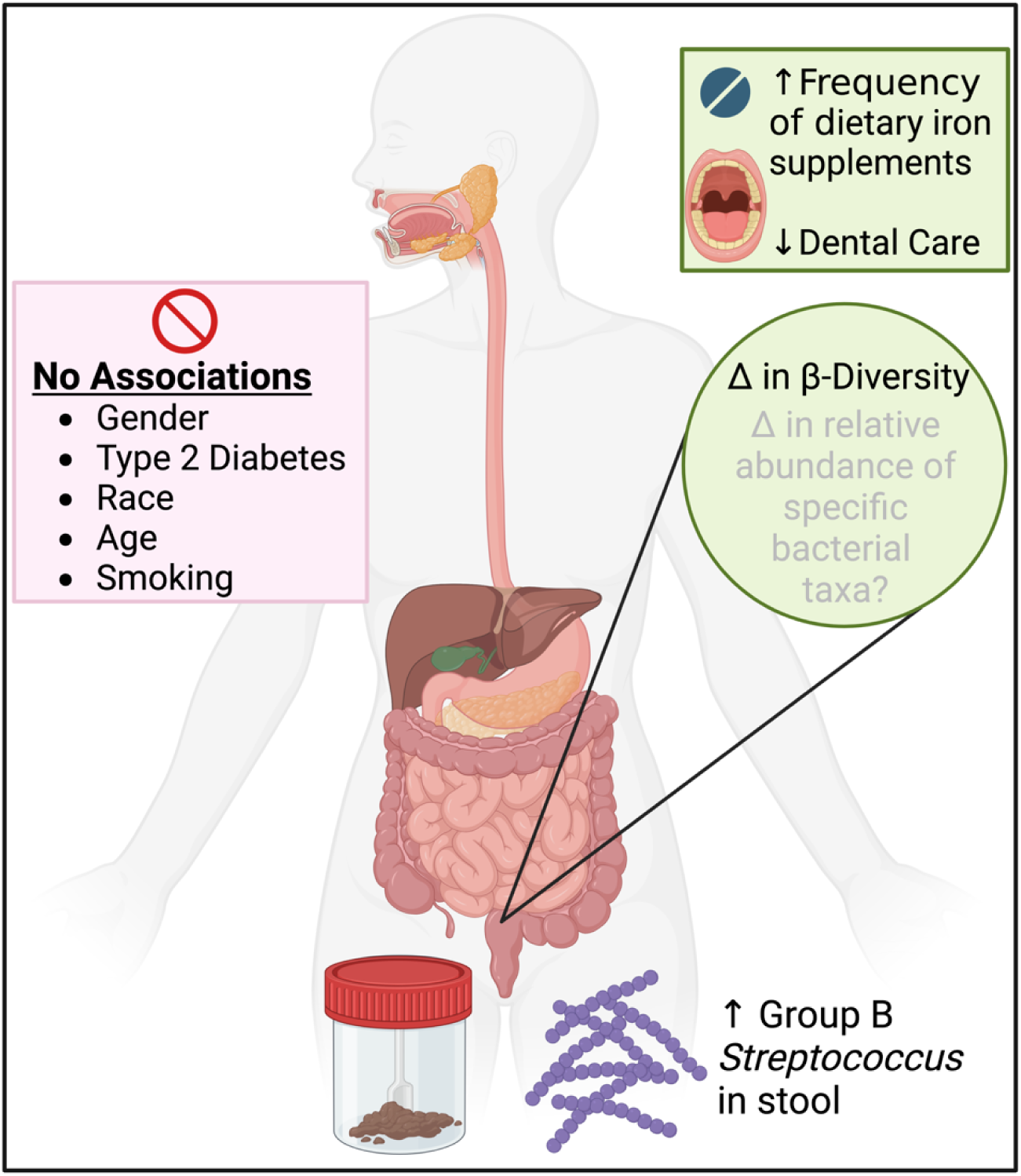
Overview of associations found with GBS in stool from study participants. No associations were found with historically known risk factors for GBS in pregnant individuals including gender, race, Type 2 diabetes, age, or smoking status. A higher burden of GBS in the stool was associated with decreased dental hygiene and an increase in frequency of dietary iron supplements. We found a change in beta-diversity of the gut microbiome was different depending on GBS carrier status. Two genera, *Ruminococcus* (p=0.0003 uncorrected, p=0.055 corrected) and *Monoglobus* (p=0.00057 uncorrected, p=0.057 corrected) trended towards being more abundant when GBS is present. Figure created with BioRender.com.

Limitations of our study include lack of longitudinal samples and using retrospective data. Since we used a biobank, we were limited to the survey questions that were asked with the initial studies and not able to add our own. Our statistical analyses demonstrate correlations and we are unable to provide causation within this study. Our final p-values were penalized for the high number of metadata variables we investigated. Strengths of our work include the large number of participant samples (754) from a large age range and backgrounds. Another strength is the extensive amount of survey data we were able to compare to GBS carrier status and abundance.

Our work provides a starting point to begin to understand risk factors for GBS carriage in the general population and in pregnant people using basic, translational, and clinical research approaches. Future work could expand on our findings including associations we found significant prior to p-value corrections. In addition, future work could use metabolomics-based approaches to identify microbiome-produced metabolites that correlate with GBS carriage. In addition, longitudinal sampling of study participants would provide important clues into the microbial and metabolic factors that influence transient carriage. Other future directions for this work include the use of experimental models to gain mechanistic insights into the associations we observe and impacts on GBS pathogenesis. This future work could include mouse models to determine the extent to which GBS colonizes distal GI and how dietary iron supplementation influence the abundance of GBS and other interacting microbiome members. Additional work could explore the extent to which transient distal GI colonization is influenced by oral GBS colonization, considering previous observations of high prevalence of oral GBS colonization [28].

## ACKNOWLEDGEMENTS

We thank Michael Liou and Courtney Van Den Elzen for statistical consulting, guidance, and constant coaching throughout this project. We thank Emmanuel Sampene and Kris Sankaran for additional statistical discussions. We thank Patrick Carney, Liz Townsend, and Shelby Sandstrom for technical support and advice with qPCR analysis. We thank the staff of the Survey of Health of Wisconsin for assistance with sample and metadata acquisition, especially Maria Nikodemova, Andy Bersch, and Hannah James.

ESC is a Medical Scientist Training Program (MSTP) student and was supported in part by MSTP grant T32GM140935 and by an NLM training grant to the Computation and Informatics in Biology and Medicine Training Program (NLM 5T15LM007359) at UW-Madison. This work was funded by a CDC Cooperative Agreement to the University of Wisconsin Prevention Research Center (1U48DP006383 to AJH), a gift from Judy and Sal Troia (AJH), and startup funding from the University of Wisconsin-Madison (AJH).

ESC and AJH conceptualized the project. ESC performed all laboratory experiments. ESC analyzed qPCR data. IZC analyzed 16S rRNA data. ESC and FO performed statistical analyses. ESC and IZC created the beta diversity figure. ESC created the remaining figures and all tables. GS, KA, and AJH provided feedback. IZC and FO provided written methods sections. ESC and AJH wrote the manuscript. All authors read, edited, and approved the manuscript prior to submission.

## Availability of data and materials

Raw data and code for qPCR, 16S rRNA amplicon gene, and univariate and multivariate modeling analysis and figure creation along with metadata used in this study are available at: https://github.com/escowley/GroupBStreptococcus_HumanGut. All 16S rRNA sequences are deposited in the NCBI SRA under BioProject ID PRJNA999362. All SHOW and WARRIOR questionnaires and surveys used to obtain metadata from participants are available upon request at https://www.med.wisc.edu/show/data-service-center/.

## Declarations

### Ethics approval and consent to participate

Human stool samples were previously collected and banked by the Survey of Health of Wisconsin from 2016-2017, which was reviewed and approved by the University of Wisconsin-Madison Institutional Review Board (Protocol #2013-0251). For this study, we used 754 of those stool samples from participants who agreed to have their samples used in future research. Our study was reviewed and approved by the University of Wisconsin-Madison Institutional Review Board (Protocol #2021-0025).

### Consent for publication

All authors have read and approved the submission of the manuscript and provide consent for publication.

## SUPPLEMENTARY TABLE LEGENDS

**Supplementary Table 1.** Random forest classification results using the microbiome composition to predict GBS carrier status.

**Supplementary Table 2.** Summary of variables included in multivariate modeling, including medians and interquartile ranges for each variable. Diet variables that are listed twice represent calculations from separate summary diet calculations. P-values denote differences between participants with GBS present and those without GBS for each variable.

**Supplementary Table 3.** Univariate associations of GBS presence and absence with metadata variables, including false discovery rate corrections.

**Supplementary Table 4.** Multivariate associations of GBS presence and absence with metadata variables, including false discovery rate corrections.

**Supplementary Table 5.** Univariate associations of GBS abundance with metadata variables, including false discovery rate corrections. Variables highlighted in green were statistically significant (p<0.05) after false discover rate correction using the Benjamini-Hochberg procedure.

**Supplementary Table 6.** Multivariate associations of GBS abundance with metadata variables, including false discovery rate corrections. Variables highlighted in green were statistically significant (p<0.05) after false discover rate correction using the Benjamini-Hochberg procedure.

**Supplementary Table 7.** Subgroup analysis (females only) of univariate associations of GBS presence and absence with metadata variables, including false discovery rate corrections.

**Supplementary Table 8.** Subgroup analysis (females only) of univariate associations of GBS abundance with metadata variables, including false discovery rate corrections. Variables highlighted in green were statistically significant (p<0.05) after false discover rate correction using the Benjamini-Hochberg procedure.

## REFERENCES

1. Schuchat A. Epidemiology of Group B Streptococcal Disease in the United States: Shifting Paradigms. Clin Microbiol Rev. 1998;11:497–513.

2. Armistead B, Oler E, Adams Waldorf K, Rajagopal L. The Double Life of Group B *Streptococcus*: Asymptomatic Colonizer and Potent Pathogen. Journal of Molecular Biology. 2019;431:2914–31.

3. Skoff TH, Farley MM, Petit S, Craig AS, Schaffner W, Gershman K, et al. Increasing burden of invasive group B streptococcal disease in nonpregnant adults, 1990-2007. Clin Infect Dis. 2009;49:85–92.

4. Schwartz B, Schuchat A, Oxtoby MJ, Cochi SL, Hightower A, Broome CV. Invasive group B streptococcal disease in adults. A population-based study in metropolitan Atlanta. JAMA. 1991;266:1112–4.

5. Farley MM, Harvey C, Stull T, Smith JD, Schuchat A, Wenger JD, et al. A Population-Based Assessment of Invasive Disease Due to Group B *Streptococcus* in Nonpregnant Adults. N Engl J Med. 1993;328:1807–11.

6. Russell NJ, Seale AC, O’Driscoll M, O’Sullivan C, Bianchi-Jassir F, Gonzalez-Guarin J, et al. Maternal Colonization With Group B *Streptococcus* and Serotype Distribution Worldwide: Systematic Review and Meta-analyses. Clinical Infectious Diseases. 2017;65:S100–11.

7. Verani JR, McGee L, Schrag SJ, Division of Bacterial Diseases, National Center for Immunization and Respiratory Diseases, Centers for Disease Control and Prevention (CDC). Prevention of perinatal group B streptococcal disease--revised guidelines from CDC, 2010. MMWR Recomm Rep. 2010;59:1–36.

8. Edwards JM, Watson N, Focht C, Wynn C, Todd CA, Walter EB, et al. Group B Streptococcus (GBS) Colonization and Disease among Pregnant Women: A Historical Cohort Study [Internet]. Infectious Diseases in Obstetrics and Gynecology. Hindawi; 2019 [cited 2021 Jan 11]. p. e5430493. Available from: https://www.hindawi.com/journals/idog/2019/5430493/

9. Krohn MA, Hillier SL, Baker CJ. Maternal Peripartum Complications Associated with Vaginal Group B *Streptococci* Colonization. The Journal of Infectious Diseases. 1999;179:1410–5.

10. Deutscher M, Lewis M, Zell ER, Taylor TH Jr, Van Beneden C, Schrag S, et al. Incidence and Severity of Invasive Streptococcus pneumoniae, Group A Streptococcus, and Group B *Streptococcus* Infections Among Pregnant and Postpartum Women. Clinical Infectious Diseases. 2011;53:114–23.

11. Seale AC, Bianchi-Jassir F, Russell NJ, Kohli-Lynch M, Tann CJ, Hall J, et al. Estimates of the Burden of Group B Streptococcal Disease Worldwide for Pregnant Women, Stillbirths, and Children. Clinical Infectious Diseases. 2017;65:S200–19.

12. Wood EG, Dillon HC. A prospective study of group B streptococcal bacteriuria in pregnancy. Am J Obstet Gynecol. 1981;140:515–20.

13. Persson K, Bjerre B, Elfström L, Polberger S, Forsgren A. Group B *Streptococci* at delivery: high count in urine increases risk for neonatal colonization. Scand J Infect Dis. 1986;18:525–31.

14. Melin P. Neonatal group B streptococcal disease: from pathogenesis to preventive strategies. Clin Microbiol Infect. 2011;17:1294–303.

15. Madrid L, Seale AC, Kohli-Lynch M, Edmond KM, Lawn JE, Heath PT, et al. Infant Group B Streptococcal Disease Incidence and Serotypes Worldwide: Systematic Review and Meta-analyses. Clin Infect Dis. 2017;65:S160–72.

16. Edmond KM, Kortsalioudaki C, Scott S, Schrag SJ, Zaidi AKM, Cousens S, et al. Group B streptococcal disease in infants aged younger than 3 months: systematic review and meta-analysis. Lancet. 2012;379:547–56.

17. Badri MS, Zawaneh S, Cruz AC, Mantilla G, Baer H, Spellacy WN, et al. Rectal Colonization with Group B *Streptococcus*: Relation to Vaginal Colonization of Pregnant Women. The Journal of Infectious Diseases. 1977;135:308–12.

18. Meyn LA, Krohn MA, Hillier SL. Rectal Colonization by Group B *Streptococcus* as a Predictor of Vaginal Colonization. Am J Obstet Gynecol. 2009;201:76.e1–76.e7.

19. Francois Watkins LK, McGee L, Schrag SJ, Beall B, Jain JH, Pondo T, et al. Epidemiology of Invasive Group B Streptococcal Infections Among Nonpregnant Adults in the United States, 2008-2016. JAMA Intern Med. 2019;179:479.

20. Langley G, Schaffner W, Farley MM, Lynfield R, Bennett NM, Reingold A, et al. Twenty Years of Active Bacterial Core Surveillance. Emerg Infect Dis. 2015;21:1520–8.

21. Phares CR. Epidemiology of Invasive Group B Streptococcal Disease in the United States, 1999-2005. JAMA. 2008;299:2056.

22. Prevention of Group B Streptococcal Early-Onset Disease in Newborns: ACOG Committee Opinion Summary, Number 797. Obstetrics & Gynecology. 2020;135:489– 92.

23. Pitts SI, Maruthur NM, Langley GE, Pondo T, Shutt KA, Hollick R, et al. Obesity, Diabetes, and the Risk of Invasive Group B Streptococcal Disease in Nonpregnant Adults in the United States. Open Forum Infectious Diseases. 2018;5:ofy030.

24. Manning SD, Neighbors K, Tallman PA, Gillespie B, Marrs CF, Borchardt SM, et al. Prevalence of Group B *Streptococcus* Colonization and Potential for Transmission by Casual Contact in Healthy Young Men and Women. Clinical Infectious Diseases. 2004;39:380–8.

25. Bliss SJ, Manning SD, Tallman P, Baker CJ, Pearlman MD, Marrs CF, et al. Group B *Streptococcus* Colonization in Male and Nonpregnant Female University Students: A Cross-Sectional Prevalence Study. Clinical Infectious Diseases. 2002;34:184–90.

26. Edwards MS, Rench MA, Palazzi DL, Baker CJ. Group B Streptococcal Colonization and Serotype-Specific Immunity in Healthy Elderly Persons. Clinical Infectious Diseases. 2005;40:352–7.

27. van Kassel MN, Janssen SWCM, Kofman S, Brouwer MC, van de Beek D, Bijlsma MW. Prevalence of group B streptococcal colonization in the healthy non-pregnant population: a systematic review and meta-analysis. Clinical Microbiology and Infection. 2021;27:968–80.

28. Martins ER, Nascimento do Ó D, Marques Costa AL, Melo-Cristino J, Ramirez M. Characteristics of Streptococcus agalactiae Colonizing Nonpregnant Adults Support the Opportunistic Nature of Invasive Infections. Microbiology Spectrum. 2022;10:e01082–22.

29. Eggers S, Malecki KM, Peppard P, Mares J, Shirley D, Shukla SK, et al. Wisconsin microbiome study, a cross-sectional investigation of dietary fibre, microbiome composition and antibiotic-resistant organisms: rationale and methods. BMJ Open. 2018;8:e019450.

30. Nieto FJ, Peppard PE, Engelman CD, McElroy JA, Galvao LW, Friedman EM, et al. The Survey of the Health of Wisconsin (SHOW), a novel infrastructure for population health research: rationale and methods. BMC Public Health. 2010;10:785.

31. Bergh K, Stoelhaug A, Loeseth K, Bevanger L. Detection of group B *Streptococci* (GBS) in vaginal swabs using real-time PCR with TaqMan probe hybridization. Indian J Med Res. 2004;119:221–3.

32. Bergseng H, Bevanger L, Rygg M, Bergh K. Real-time PCR targeting the sip gene for detection of group B *Streptococcus* colonization in pregnant women at delivery. Journal of Medical Microbiology. 2007;56:223–8.

33. Escobar DF, Diaz-Dinamarca DA, Hernández CF, Soto DA, Manzo RA, Alarcón PI, et al. Development and analytical validation of real-time PCR for the detection of *Streptococcus agalactiae* in pregnant women. BMC Pregnancy Childbirth [Internet]. 2020 [cited 2020 Dec 14];20. Available from: https://www.ncbi.nlm.nih.gov/pmc/articles/PMC7285471/

34. Carrillo-Ávila JA, Gutiérrez-Fernández J, González-Espín AI, García-Triviño E, Giménez-Lirola LG. Comparison of qPCR and culture methods for group B *Streptococcus* colonization detection in pregnant women: evaluation of a new qPCR assay. BMC Infectious Diseases. 2018;18:305.

35. Khatami A, Randis TM, Chamby A, Hooven TA, Gegick M, Suzman E, et al. Improving the Sensitivity of Real-time PCR Detection of Group B *Streptococcus* Using Consensus Sequence-Derived Oligonucleotides. Open Forum Infect Dis [Internet]. 2018 [cited 2021 Mar 3];5. Available from: https://www.ncbi.nlm.nih.gov/pmc/articles/PMC6051451/

36. Bolyen E, Rideout JR, Dillon MR, Bokulich NA, Abnet CC, Al-Ghalith GA, et al. Reproducible, interactive, scalable and extensible microbiome data science using QIIME 2. Nat Biotechnol. 2019;37:852–7.

37. Callahan BJ, McMurdie PJ, Rosen MJ, Han AW, Johnson AJA, Holmes SP. DADA2: High-resolution sample inference from Illumina amplicon data. Nature Methods. 2016;13:581–3.

38. Katoh K, Misawa K, Kuma KI, Miyata T. MAFFT: A novel method for rapid multiple sequence alignment based on fast Fourier transform. Nucleic Acids Research. 2002;30:3059–66.

39. Bokulich NA, Kaehler BD, Rideout JR, Dillon M, Bolyen E, Knight R, et al. Optimizing taxonomic classification of marker-gene amplicon sequences with QIIME 2’s q2-feature-classifier plugin. Microbiome. 2018;6:1–17.

40. Quast C, Pruesse E, Yilmaz P, Gerken J, Schweer T, Yarza P, et al. The SILVA ribosomal RNA gene database project: Improved data processing and web-based tools. Nucleic Acids Research. 2013;41:590–6.

41. McMurdie PJ, Holmes S, Kindt R, Legendre P, O’Hara R. phyloseq: An R Package for Reproducible Interactive Analysis and Graphics of Microbiome Census Data. Watson M, editor. PLoS ONE. 2013;8:e61217.

42. Davis NM, Proctor DiM, Holmes SP, Relman DA, Callahan BJ. Simple statistical identification and removal of contaminant sequences in marker-gene and metagenomics data. Microbiome. 2018;6:1–14.

43. Lin H, Peddada S Das. Analysis of compositions of microbiomes with bias correction. Nature Communications. 2020;11:1–11.

44. Segata N, Izard J, Waldron L, Gevers D, Miropolsky L, Garrett WS, et al. Metagenomic biomarker discovery and explanation. Genome Biology. 2011;12:R60.

45. Cao Y, Dong Q, Wang D, Zhang P, Liu Y, Niu C. microbiomeMarker: an R/Bioconductor package for microbiome marker identification and visualization. Bioinformatics. 2022;38:4027–9.

46. Tang ZZ, Chen G, Alekseyenko A V., Li H. A general framework for association analysis of microbial communities on a taxonomic tree. Bioinformatics. 2017;33:1278– 85.

47. Liaw A, Wiener M. Classification and Regression by randomForest. R News. 2002;2:18–22.

48. Kwatra G, Adrian PV, Shiri T, Buchmann EJ, Cutland CL, Madhi SA. Serotype-Specific Acquisition and Loss of Group B *Streptococcus* Recto-Vaginal Colonization in Late Pregnancy. PLOS ONE. 2014;9:e98778.

49. Abranches J, Zeng L, Kajfasz JK, Palmer SR, Chakraborty B, Wen ZT, et al. Biology of Oral *Streptococci*. Microbiology Spectrum. 2018;6:10.1128/microbiolspec.gpp3-0042–2018.

50. Hillman ET, Lu H, Yao T, Nakatsu CH. Microbial Ecology along the Gastrointestinal Tract. Microbes and environments. 2017;32:300–13.

51. Cassat JE, Skaar EP. Iron in Infection and Immunity. Cell Host & Microbe. 2013;13:509–19.

52. Clancy A, Loar JW, Speziali CD, Oberg M, Heinrichs DE, Rubens CE. Evidence for siderophore-dependent iron acquisition in group B *Streptococcus*. Molecular Microbiology. 2006;59:707–21.

53. Yilmaz B, Li H. Gut Microbiota and Iron: The Crucial Actors in Health and Disease. Pharmaceuticals (Basel). 2018;11:98.

54. Rusu IG, Suharoschi R, Vodnar DC, Pop CR, Socaci SA, Vulturar R, et al. Iron Supplementation Influence on the Gut Microbiota and Probiotic Intake Effect in Iron Deficiency—A Literature-Based Review. Nutrients. 2020;12:1993.

55. Dahesh S, Hensler ME, Van Sorge NM, Gertz RE, Schrag S, Nizet V, et al. Point Mutation in the Group B Streptococcal pbp2x Gene Conferring Decreased Susceptibility to β-Lactam Antibiotics. Antimicrobial Agents and Chemotherapy. 2008;52:2915–8.

56. Longtin J, Vermeiren C, Shahinas D, Tamber GS, McGeer A, Low DE, et al. Novel Mutations in a Patient Isolate of *Streptococcus agalactiae* with Reduced Penicillin Susceptibility Emerging after Long-Term Oral Suppressive Therapy. Antimicrobial Agents and Chemotherapy. 2011;55:2983–5.

57. Kimura K, Suzuki S, Wachino J, Kurokawa H, Yamane K, Shibata N, et al. First molecular characterization of group B *Streptococci* with reduced penicillin susceptibility. Antimicrob Agents Chemother. 2008;52:2890–7.

58. Persson E, Berg S, Bergseng H, Bergh K, Valsö-lyng R, Trollfors B. Antimicrobial susceptibility of invasive group B streptococcal isolates from south-west Sweden 1988– 2001. Scandinavian Journal of Infectious Diseases. 2008;40:308–13.

59. Simoes JA, Aroutcheva AA, Heimler I, Faro S. Antibiotic Resistance Patterns of Group B Streptococcal Clinical Isolates. Infectious Diseases in Obstetrics and Gynecology. NaN/NaN/NaNxs;12:1–8.

60. Dethlefsen L, Huse S, Sogin ML, Relman DA. The Pervasive Effects of an Antibiotic on the Human Gut Microbiota, as Revealed by Deep 16S rRNA Sequencing. PLOS Biology. 2008;6:e280.

